# Searching for Principles of Brain Computation

**DOI:** 10.1101/094102

**Authors:** Wolfgang Maass

## Abstract

**Highlights:**

- Hints for computational principles from experimental data
- Computational role of diverse network components
- Emergence and computational role of assemblies
- Probabilistic inference through stochastic network dynamics
- Ongoing network rewiring and compensation through synaptic sampling

**Abstract:** Experimental methods in neuroscience, such as calcium-imaging and recordings with multielectrode arrays, are advancing at a rapid pace. They produce insight into the simultaneous activity of large numbers of neurons, and into plasticity processes in the brains of awake and behaving animals. These new data constrain models for neural computation and network plasticity that underlie perception, cognition, behavior, and learning. I will discuss in this short article four such constraints: Inherent recurrent network activity and heterogeneous dynamic properties of neurons and synapses, stereotypical spatio-temporal activity patterns in networks of neurons, high trial-to-trial variability of network responses, and functional stability in spite of permanently ongoing changes in the network. I am proposing that these constraints provide hints to underlying principles of brain computation and learning.

## Constraint/Principle 1: Neural circuits are highly recurrent networks consisting of different types of neurons and synapses with diverse dynamic properties

Most computations in our current generation of digital computers have a feedforward organization, where specific network modules carry out specific subcomputations and transmit their results to the next module. One advantage of this computational organization is that it is easy to understand and control. Also deep learning networks favor feedforward computation (and backwards propagation of errors or predictions) because this organization best supports currently known algorithms for network learning. But nature has apparently discovered a way of fully using recurrent neural networks for reliable computation and learning that is based on different principles. Already evolutionary very old nervous systems, such as those of hydra [1], C-elegans [2], and zebrafish [3] are highly recurrent and exhibit a complex global network dynamics, similar as brain networks in more advanced species.

These activity patterns differ in several aspects from those that we encounter in our digital computers. Hence recent reviews [4], [5] have emphasized the need to understand the basic principles of brain computations in recurrent networks of neurons.

Numerous theoretical and modelling studies have analyzed the dynamics of simple recurrent networks of homogeneous types of neurons and synapses, see e.g. [6], [7], [8]. But neural networks in the brain consist of different types of neurons [9] that are connected by different types of synapses with heterogeneous shortterm and longterm dynamics [10], [11], [12]. These features of biological networks of neurons make a theoretical analysis difficult, and they constrain computational models. In particular, biological networks of neurons are not well-suited for emulating generic computations of Boolean circuits or artificial neural networks. For example, it would be difficult to implement tensor product variable binding [13] in a neural network model which takes into account that there are excitatory and inhibitory neurons with different dynamic properties, that neurons do not emit analog values but spikes at low firing rates, and that synapses are subject to noise and shortterm dynamics (i.e., mixtures of paired pulse facilitation and depression). The shortterm dynamics of synapses lets the amplitudes of postsynaptic potentials decrease or increase for a sequence of spikes in dependence of the pattern of preceding spikes. This history-dependence obstructs a stable transmission of spikes and firing rates, which we would need for emulating a Boolean circuit or artificial neural network. The obvious question is of course whether the experimentally found diversity of units, mechanisms, and time-constants in brain networks is detrimental for all types of computations, or whether it could enhance specific computational operations that nature has discovered.

One computational model for which a diversity of computational units and time constants is not detrimental, and in fact benefitial, is the liquid computing paradigm [14], [15], [16]. It is sometimes subsumed together with the somewhat similar echo-state model of [17] under the name reservoir computing, see ch. 20 of [8]. A common feature of both types of reservoir computing models is that they conceptually divide neural network computations into two stages (see Fig. 1a), a fixed generic nonlinear preprocessing stage and a subsequent stage of linear readouts that are trained for specific computational tasks. A structural difference between the liquid computing model and the echo-state model is that the latter assumes that there is no noise in the network, and that analog outputs of computational units can be transmitted with arbitrary precision to other units. In contrast, the liquid computing model is geared towards biological neural networks, where noise, diversity of units, and temporal aspects related to spikes play a prominent role.

A diversity of units and time constants in a recurrent neural network causes no problem if one analyzes its computational contribution from the perspective of a projection- or readout neuron (such as pyramidal cells on layers 2/3 and 5 [9]) that receives synaptic inputs from thousands of neurons within the recurrent neural network, and extracts information for other networks. From this perspective it is not required that neurons in the recurrent neural network complete specific subcomputations. It suffices if diverse neurons and subcircuits within the recurrent network produce a large number of potentially useful features and nonlinear combinations of such features, out if which a projection neuron can select and combine through a weighted sum — or a more complex dendritic integration — useful information for its target networks. In this way even a seemingly chaotic dynamics of a recurrent local network can make a useful computational contribution [14],[15],[18],[16],[19],[20].

This perspective raises the question how a recurrent neural network could optimally support through generic computational preprocessing subsequent readout neurons. Some theoretical foundation (see [15], [18], [21], [22], and Fig. 1 a for details) arises through a link to one of the most successful learning approaches in machine learning: Support Vector Machines (SVMs; [23]). A SVM also consists of two stages: a generic nonlinear preprocessing stage (called kernel) and linear readouts. The kernel projects external input vectors x_1_, x_2_,… nonlinearly onto vectors h_1_, h_2_,…. in a much higher dimensional space. One can view a large nonlinear recurrent neural network as an implementation of such a kernel, where the network response hi to an input x_i_ corresponds to the kernel output. This network response h_i_ can be defined for example as the high-dimensional vector that records for each neuron in the network its recent firing activity, say within the last 30 ms (as in Fig. 3b). This network response h_i_ provides then the synaptic input to any readout neurons. It represents the "visible” part of the network state, while other "hidden” dimensions of the true network state, such as the current internal state of dynamic synapses is not visible for readout neurons [16]. If the map from x_i_ to h_i_ is nonlinear, the network can increase the expressive capability of subsequent linear readouts. For example, if the network states h_1_, h_2_,… would contain all products of components of the network inputs x_1_, x_2_,…, a linear readout from these network states attains the same expressive capability as a quadratic readout function in terms of the original network inputs x_1_, x_2_,…. The quality of the kernel operation of a neural network can be measured by the dimension of the linear vector space that is spanned by the ensemble of network states h_1_, h_2_,… which result for some finite ensemble of different network inputs x_1_, x_2_,…. This dimension is equal to the rank of the matrix with columns h_1_, h_2_,…. This approachto measure the computational power of a neural circuit through a dimensionality analysis was introduced in [21], [18], and later applied to experimental data in [22], see [24] for a review. It provides an alternative to approaches based on the analysis of neural codes and tuning curves of individual neurons for specific simple stimuli. It provides instead a paradigm for analyzing neural codes for complex natural stimuli x_i_ on the network level – from the perspective of neural readouts. The kernel property of a neural network would be theoretically optimal if it would map any ensemble of different external inputs x_1_, x_2_,… onto linearly independent network states h_1_, h_2_,…. A linear readout can assign through a proper choice of its weights any desired output values o_1_, o_2_,… to linearly independent network states h_1_, h_2_,…. If the vectors h_1_, h_2_,… are not linearly independent, then the rank of the matrix with these columns tells us how much of these theoretically ideal expressive capabilities of linear readouts remain. A more subtle analysis is needed to integrate noise-tolerance into this network level analysis of neural coding [21], [18], [22], since a readout needs to be able to assign target outputs in a trial-invariant manner. Hence one needs to distinguish linear independence of network states h_i_ caused by saliently different inputs x_i_ from accidental linear independence caused by noise. But if the network is sufficiently large and nonlinear, it tends to endow a simple linear readout even in the presence of noise with the computational and learning capability of a more complex nonlinear readout. In addition, the resulting two stage network has a very desirable learning dynamics if only the weights of a linear readout are adjusted: there are no local minima in its error function – hence gradient descent arrives in the end at the global optimum.

One other benefit of such a two-stage computational model is its multiplexing efficiency: The same first stage (the kernel) can be shared by an unlimited number of subsequent linear projection neurons (indicated on the right in Fig. 1a), that learn to extract different computational results for their specific target networks (see a simple demo in Fig. 1b). This feature does not depend on specific aspects of the model, but is shared with any model wich proposes that generic cortical microcircuits generate a menu of features that supports a variety of downstream computations. Such multiplexing of computations through parallel readouts from a common recurrent network provides an alternative to models based on a precise ice-cube-like spatial organization of sub-computations in a cortical column, see e.g. [25] for a discussion. Recent experimental data [26] suggest that different projection neurons do in fact extract from the same local microcircuit quite different results.

So far I have only addressed static computations on batch input vectors x_i_. A further computational benefit of having diverse units in a neural network, especially units with a wide spread of time constants, becomes apparent if one takes into account that many brain computations have to integrate information from several preceding time windows.An important class of such computations are computations on time series with a fading memory. These are computations where the output at time t may also depend on inputs that have arrived before time t in the recent past. Surprisingly, even the arguably most complex nonlinear transformations that map input time series onto output time series with a fading memory, Volterra series, can be implemented through simple memory-less readouts from any ensemble of filters that have a sufficiently wide spread of time constants. If one views synapses with their inherent shortterm dynamics as filters, then it suffices if the network contains synapses with diverse shortterm dynamics. More precisely the following separation property of the ensemble of filters is relevant (see Theorem 1 in [14] and [27] for further details): Does at least one of the filters produce at the current time t different output values for two input time series that differed at some point in the recent past? It has recently been shown that both cultured neural circuits [28] and ganglion cells in the retina [29] have a good separation property of this kind. A good separation property entails that output o_3_ of a linear readout at time step 3 may depend nonlinearly not only on the current network input x_3_, but also on preceding network inputs x_1_ and x_2_. The liquid computing model (see Fig. 1a) postulates that the separation property is, in addition to the previously discussed kernel property, a basic computational property of generic neural circuits. This prediction of the model was subsequently verified through recordings from visual [30] and auditory [31] primary cortex. In principle, even a working memory can be composed according to this analyzis from local units or modules of a recurrent neural network that have different time constants [32], [33], [34].

An interesting question is which details of biological neural circuits are essential for maximizing their kernel-and separation property (see [15], [35] for some first results). Recurrent connections, diversity of neuron types, and diversity of synapse types all appear to contribute to the kernel-and separation property. But not all of these features appear to be necessary for that. An alternative view of the experimentally found complexity of neural circuits is that tightly structured connectivity, homogeneity of neurons, and homogeneity of synapses are essential properties of human-designed computational circuits, but are task-irrelevant dimensions for biological neural circuits, because readout neurons with adaptive capabilities can compensate for inhomogeneities and deficiencies of the circuits which provide inputs to them. In other words many details of neural circuits can be viewed as task-irrelevant dimensions. A more specific functional role of diverse types of synaptic dynamics for stabilizing network activity was proposed in [36].

The importance of readouts becomes even larger if one no longer assumes that their output only affects downstream networks. Mathematical results [37] imply that the capability of the liquid computing model is substantially enhanced if linear readout neurons — that are trained for specific tasks — are allowed to project their output also back into the local network (see dashed loop at the bottom of Fig 1a). In fact, most projection neurons from a generic cortical microcircuit do have axon collaterals, which carry out such back projections. The essential structural difference to the model without feedback is that now the training also affects the dynamics of the recurrent network itself. Under ideal conditions without noise this model with feedback acquires the computational power of a universal Turing machine [37]. But computer simulations (see Fig. 1c, d) show that the feedback also adds in the presence of noise important computational capabilities to the liquid computing model: It now can remember salient inputs in its internal state for unlimited time (see Fig.1 c, d). Furthermore it can switch its computational function in dependence of its internal state, and hence also in dependence of external cues (Fig. 1e). Experimental data [38] have subsequently shown that cortical networks of neurons do in fact have these theoretically predicted enhanced computational capabilities.

**Figure 1:**
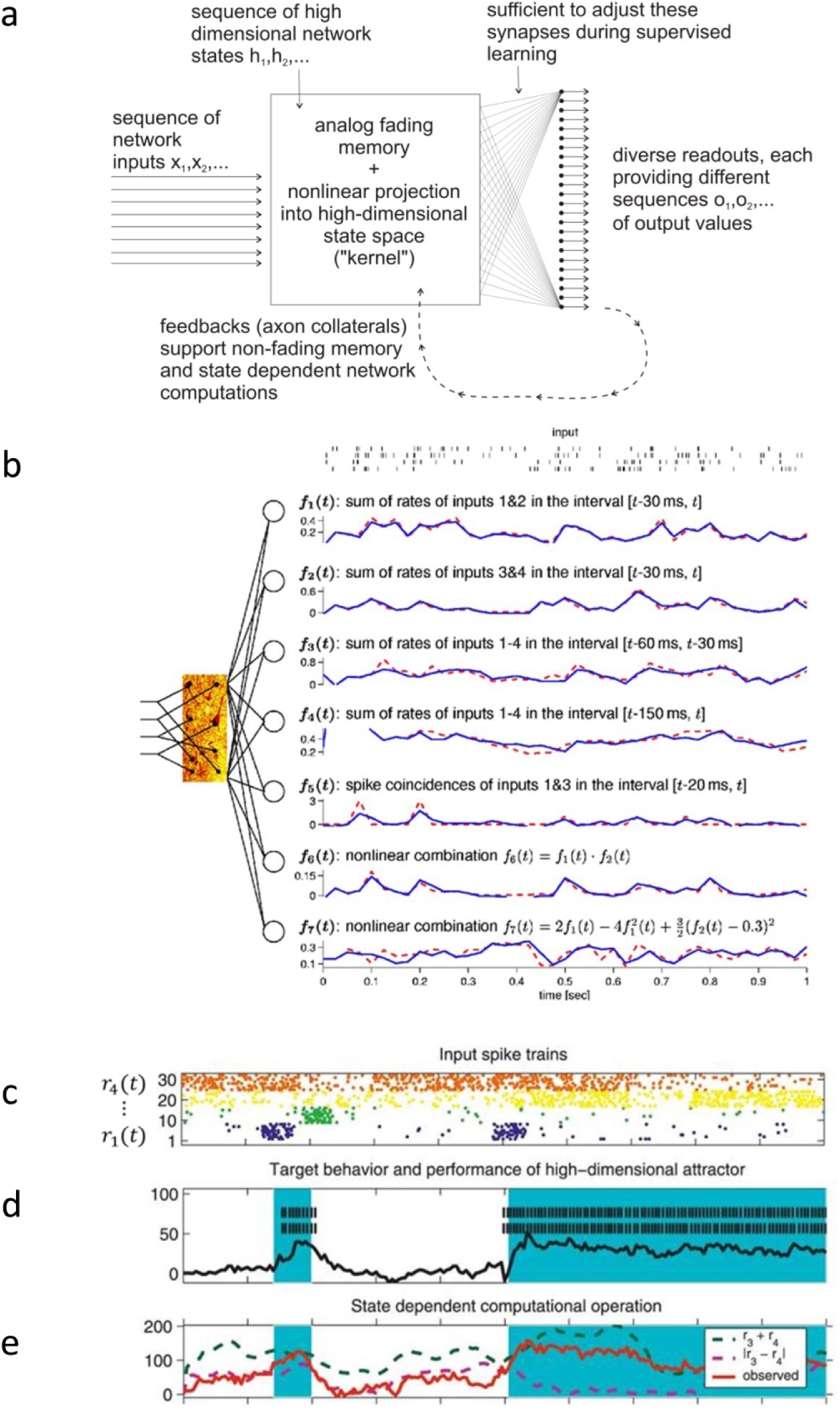
Computational paradigms resulting from principle 1. a) Generic computational model; b) Demonstration that a randomly connected network of 135 spiking neurons with diverse shortterm plasticity of synaptic connections supports multiplexing: Different linear readouts can be trained to produce simultaneously different online computations, in this example on two time-varying firing rates f_1_(t) and f_2_(t) represented by the spiking activity of the first 2 and the last 2 input neurons shown at the top [39]. Outputs of the linear readouts are plotted as blue curves, target outputs as dashed red curves. c) — e): Demonstration of additional computational properties that arise with feedback from trained readouts: nonfading memory and context-depending switching of computations. c): 4 spike input streams with timevarying Poisson firing rates r_1_(t),…, r_4_(t). d): Two spiking readouts with feedback (their spike outputs are shown in black) were trained to remember which of the first two input streams had last exhibited a burst (time points where r_1_(t) had the most recent burst are marked in blue). e): Another readout was trained to output spikes with rates r_3_(t) + r_4_(t) or |r_3_(t) — r_4_(t)| (both shown as dashed curves) in dependence of the binary state represented in d). The orange curve shows the resulting output of this readout neuron, that approximates r_3_(t) + r_4_(t) during one network state (indicated by blue) and otherwise |r_3_(t) + r_4_(t)|. Fig 1b is reprinted from "Theory of the Computational Function of Microcircuit Dynamics,” in "Microcircuits: The Interface between Neurons and Global Brain Function," edited by Sten Grillner and Ann M. Garybiel, published by "The MIT Press, pp. 371-390, 2006”, with kind permission from The MIT Press. Fig 1d-e is reprinted from [37] "Computational aspects of feedback in neural circuits”, in "PLOS Computational Biology, 3(1):e165, 2007”, with kind permission from The PLOS Journals.

Training of linear readouts with feedback requires well-tuned learning methods because the resulting closed loop tends to amplify the impact of changes of synaptic weights to readout neurons. Two successful methods for supervised learning [37], [19] employ and refine related methods for echo state networks [40]. These learning methods work well in simulations, but do not aspire to be biologically realistic. Biologically more realistic learning methods based on reinforcement learning were proposed in [20] and [41].

## Constraint/Principle 2: Neural network activity is dominated by variations of assembly activations

From the perspective of some theories of neural coding and computations it would be desirable that different neurons in a local network can encode independently of each other specific features of a sensory stimulus. However virtually all simultaneous recordings from many neurons within a local patch of cortex suggest that the joint activity patterns of nearby neurons are restricted to variations of a rather small repertoire of spatio-temporal firing patterns. One usually refers to these patterns as assemblies, assembly sequences, or packets of activity [42]. It was shown in [43] that patches of auditory cortex typically respond with variations of just one or two different joint activity patterns to a repertoire of over 60 auditory stimuli, and to continuously morphed stimuli. Also patches of V1 in rodents appear to respond to natural movies with variations of just a few joint activity patterns [44]. Furthermore a small repertoire of activity patterns tends to occur also spontaneously [45]. The fact that a small repertoire of joint activity patterns also occurs in slice [46] supports the conjecture that these patterns are consequences of network architecture and parameters that result from an interplay of the genetic code and plasticity processes. In particular, learned behaviours have been shown to become encoded by similar stereotypical joint activity pattern in the higher cortical areas PFC (prefrontal cortex) [47] and PPC (posterior parietal cortex) [48].

However it has remained open how neural networks compute with these stereotypical joint activity patterns. In order to test this in models, one first has to find ways of inducing their emergence. [49] showed that stereotypical patterns emerge through STDP (spike-timing dependent plasticity)) in recurrent networks with very little noise even in the absence of external inputs. More recently such patterns have also been induced through STDP in such a way that they encode the class to which an input pattern belongs [50]. Whereas this model used simplified lateral inhibition, Fig. 2 a, b shows that similar pattern emerge through STDP in networks with explicitly modeled inhibitory neurons [51]. Furthermore it has been shown that input-dependent assemblies also emerge in models that employ in addition synaptic plasticity for inhibitory synapses [52]. One computational benefit that is suggested by these models is that assembly coding facilitates the learning task of readout neurons: They are able to learn very fast – even without supervision (see Fig. 2 c-d) – to report which assembly is currently active, and hence to which class an input pattern belongs.

Assemblies and assembly sequences had already been postulated by [53] to be tokens of network dynamics that create links between the fast time scale of spikes and the slower time scale of cognition and behaviour. [54] proposed to view assemblies as word-like codes for salient objects, concepts etc that are combined in the brain through a yet unknown type of "neural syntax”.

**Figure 2:**
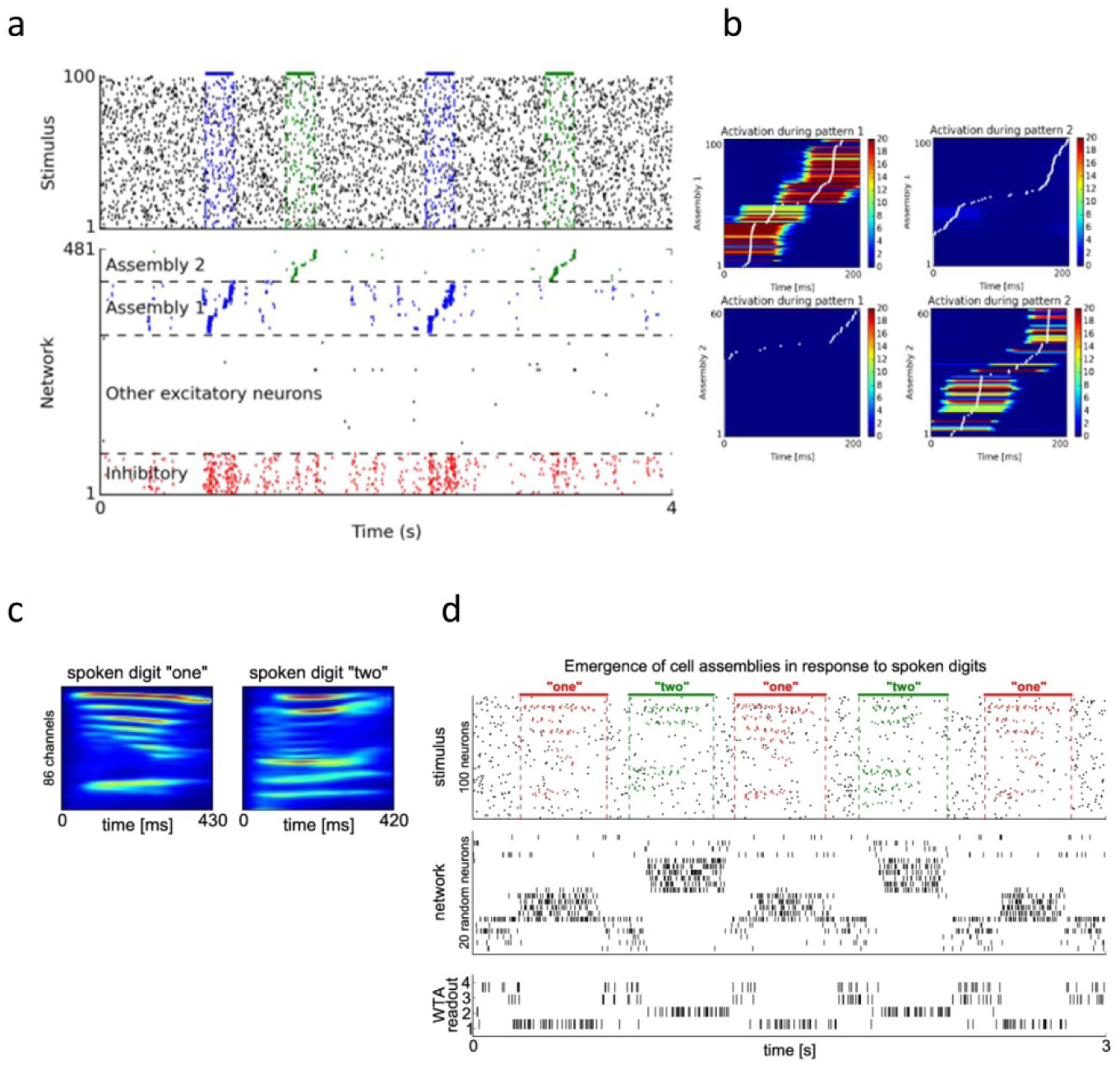
Emergence and computational use of assembly codes. a) Emergence of input-specific assemblies (or more precisely, assembly sequences) through STDP in response to repeating external input patterns (there are blue and green spike patterns which are superimposed by noise spikes shown in black). This occurs even if the input patterns (frozen Poisson patterns) have exactly the same rates and statistics as the noise input between patterns. The assembly sequences shown in a) and b) emerge after about 100 occurrences of each input pattern in a generic recurrent neural network [51]. b) The mean firing time of each excitatory neurons is marked by a white dot, with a histogram of all firing times represented through color coding in the same row (analogously as in [45]). c) Sample utterings of two spoken digits that were transformed into spike inputs shown in the top row of d) in a network simulation from [50]. The middle row of d) shows the two emergent assemblies for the two spoken words “one” and “two”. The bottom row of d) shows the firing response of 4 linear readout neurons in a WTA (winner take all) circuit. This WTA readout learns without any supervision to report the occurence of one of the two assembly sequences, and hence spoken digit classification, through the firing of neurons 1 and 2. Panels c and d of Fig. 2 are reprinted from [50] “Emergence of dynamic memory traces in cortical microcircuit models through STDP” published by “The Journal of Neuroscience, 33(28): 11515–11529”, 2013, with kind permission from The Journal of Neuroscience.

## Constraint/Principle 3: Networks of neurons in the brain are spontaneously active and exhibit high trial-to-trial variability

Virtually all neural recordings show that network responses vary substantially from trial to trial. This is not surprising, since channel kinetics in dendrites and synaptic transmission are reported to be highly stochastic [55], [56]. These data force us to add a substantial amount of variability or noise to the set of constraints for neural network computations. Again, a key question is whether this constraint can also be viewed as a principle that provides a clue for understanding the organization of brain computations. Usually noise is just seen as a nuisance in a computational system [57].

Hints for a possible benefitial role of large trial-to-trial variability for brain computations is provided by experimental data which suggest that ambiguous sensory stimuli are represented in brain networks through flickering between different network states, that each represent one possible interpretation of the ambiguous stimulus [58], [59]. Also the values of possible choices appear to be represented in monkey orbitofrontal cortex (OFC) before decision making through flickering between corresponding network states [60] on a small time scale like in Fig. 3b. These new data from simultaneous recordings from many neurons with high temporal precision suggest that “subjective decision-making involves the OFC network transitioning through multiple states, dynamically representing the value of both chosen and unchosen options” [60]. A well known approach for probabilistic inference (Markov chain Monte Carlo or MCMC sampling, see [60], [23]) suggests to interpret these experimental data as probabilistic inference through stochastic computation, more precisely through sampling from some internally stored probability distribution of network states.

Insight into the nature of such internally stored probability distribution can in principle be gained by analyzing the statistics of network states, defined for example by a binary vector with a “1” for every neuron that fires within some small time bin [61] (see Fig. 3b). The term “neural sampling” had been coined in [62] for the resulting theory of probabilistic inference through sampling in stochastically firing recurrent networks of neurons. Each neuron *v_i_* represents in this model a binary random variable *z_i_* through spikes: a spike sets the value of this random variable to 1 for some short period of time. It was shown in [62] that if synaptic weights are symmetric, a network of simple models for spiking neurons can represent the same probability distribution as a Boltzmann machine with the same architecture, although it uses a different sampling strategy. This is interesting because a Boltzmann machine is one of the most studied neural network models in machine learning for probabilistic inference and learning, and it is known that it can learn and represent any multivariate distribution over binary random variables with at most 2^nd^ order dependencies. In addition it was shown that a suitable architecture enables networks of spiking neurons with asymmetric weights to go beyond that: a spiking network can represent [63] and learn [64] any distribution over discrete variables, even with higher order dependencies as they occur for example in the explaining-away effect of visual perception [65].

Also data-based models for generic cortical microcircuits (Fig.3a) with stochastically firing neurons can carry out probabilistic inference through sampling. For example they can estimate through sampling posterior marginals such as *p*(*z*_1_|**e**) = *∑a*_2_,…,*a_m_ p*(*z*_1_,*a*_2_,–-,*a_m_*|**e**), where **e** is some external input [66]. The current external input **e** could represent for example sensory evidence and internal goals. The variables for *a_i_* for i>1 run in this formula over all possible values of random variables *z_i_* that are irrelevant for the current probabilistic inference task. The sum indicates that these variables are marginalized out, which is in general a computationally very demanding (in fact: NP-hard) operation. The binary variable *z*_1_ could represent for example the choice between two decisions, so that an estimation of the posterior marginal *p*(*z*_1_|**e**) supports Bayes-optimal decision making. The key point is that this computationally very difficult posterior marginal can be estimated quite easily through sampling: It is represented by the firing rate of the neuron *v*_1_ that corresponds to the binary random variable *z*_1_ [66]. Also sampling-based representations of time-varying probabilities–where each random variable is represented through several spiking neurons – have been examined [67], [68]. At the current time point it is not yet clear to what extent brains make use of the option to carry out probabilistic inference through sampling. To answer this question one needs further experimental insight into the relation between flickering internal states of brain networks on one hand and perception and behavior on the other hand. [58] and [60] have demonstrated that this is in principle feasible.

**Figure 3:**
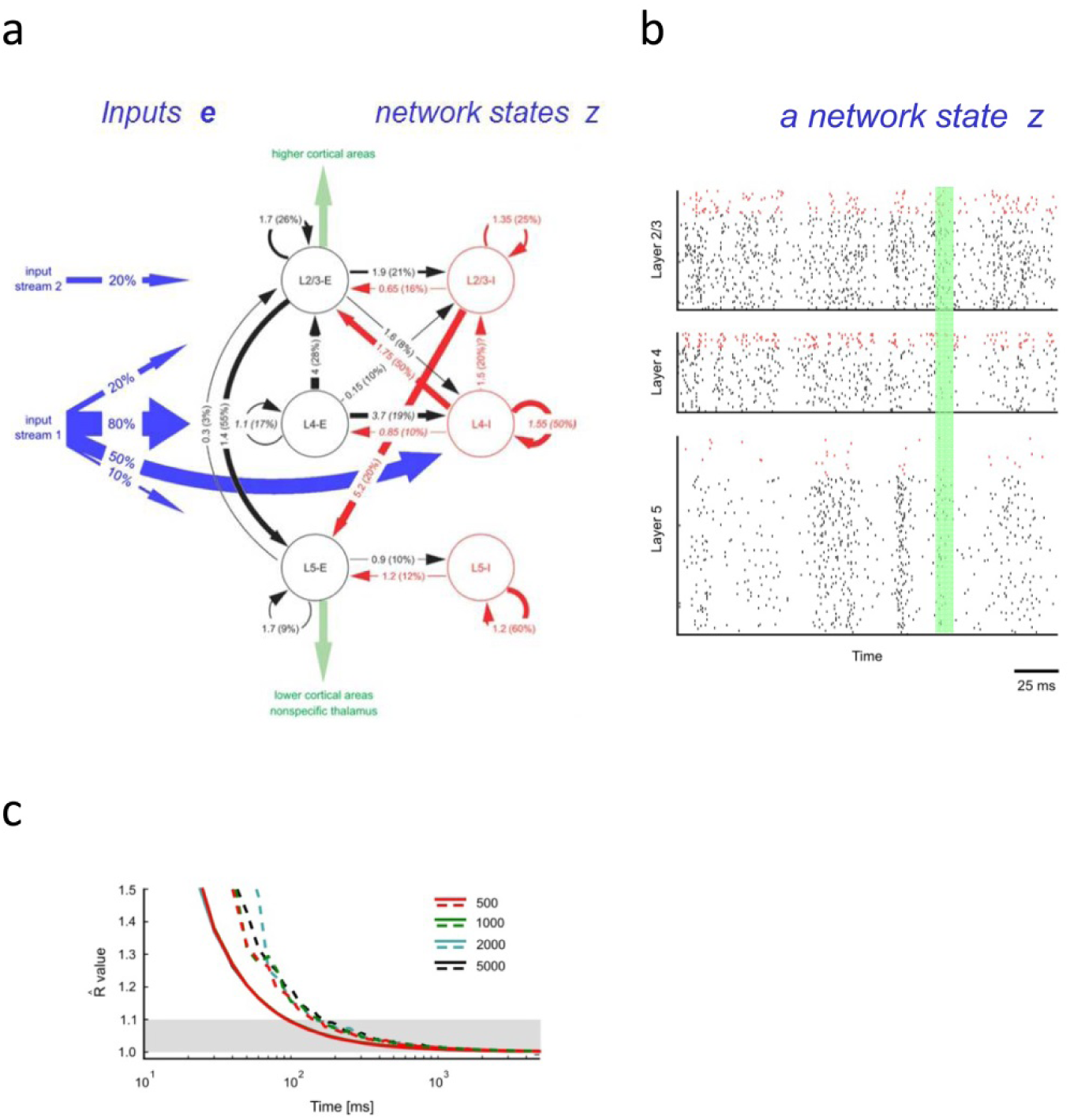
Model for probabilistic inference through sampling in a generic cortical microcircuit model with stochastically spiking neurons [66]. a) Synaptic weights and other parameters encode a unique stationary distribution p(**z**) of network states **z**. b) The network state **z** at time t can be defined for example as a binary vector [61] that records which neuron fires in a small time window around t (shaded in green). The stochastic dynamics of the network for some external input **e** can be interpreted as sampling from the conditional distribution p(**z**| **e**). c) Instead of traditional measures for computation time, the time needed to converge from an initial state to the stationary distribution of network states (not to any particular state!) becomes relevant. A standard heuristic estimate (Gelman-Rubin analysis) suggests that this convergence is quite fast – around 100ms – and independent of the network size (color coded for network sizes between 500 and 5000 neurons) for the data-based model from a). Likely reasons for independence from network size are small synaptic weights, weight normalization, and a large amount of stochasticity in the model. The Gelman-Rubin analysis suggests that convergence to the stationary distribution has taken place by the time when the curves (solid lines: mean; dashed lines: worst case) enter the gray zone below 1.1. Fig. 3 is reprinted from "Stochastic computations in cortical microcircuit models” published by "PLOS Computational Biology, 9(11):e1003311, 2013”, with kind permission from The PLOS Journals.

Stochasticity of spiking neurons conveys another computational benefit to a network: it enables the network to solve problems – for example constraint satisfaction problems – in a heuristic manner [69], [66], [57]. Here each network state(defined like in Fig. 3b) represents a possible solution to a problem, and the frequency of being in this network state encodes the quality (fitness) of the solution. This computational model is consistent with the data from [60], where easier choices were associated with fewer switches between the neural representation of the two options. The computation time for solving a task depends in such a sampling-based model on the time that the network needs until it produces, starting at some given initial state, samples from the stationary distribution of network states (see Fig. 3c), which is defined by the architecture and parameters of the network [66]. A substantial level of noise in the network and not too large synaptic weights support in general fast convergence [66].

The hypothesis that the human brain encodes substantial amounts of knowledge in the form of probabilities and probability distributions had previously been proposed in cognitive science [70], [71], [72]. Probabilistic computations have started to play a prominent role in many models in neuroscience, for example in models for multisensory integration [73] and confidence [74].

## Constraint/Principle 4: Networks of neurons in the brain provide stable computational function in spite of ongoing rewiring and network perturbations

Experimental data show that network connectivity [75], [76], [77], neurotransmitters [78] and neural codes [79] are subject to continuously ongoing changes, even in the adult brain. This constraint suggests to consider the hypothesis that the brain samples not only network states on the fast time scale of spiking activity as discussed under principle 3, but simultaneously also different network configurations on the slower time scale of network plasticity and spine dynamics (i.e., hours and days). This slower sampling of network configurations has been called synaptic sampling [80]. The synaptic sampling model suggests that brain networks do not converge to a desirable network configuration and stay there, but rather sample continuously – but at different speeds (“temperatures”) – from a posterior distribution of network configurations (Fig. 4).

Learning a posterior distribution of network configurations, rather than a specific network configuration, has been proposed to be a more attractive goal for network plasticity – for example because of better generalization capability [81]. The question how a biological network of neurons could represent and learn such a posterior distribution was described in [73] as a key open problem. [80] proposes that this posterior distribution is represented by a stationary distribution of network configurations in the Markov chain that is defined by the stochastic dynamics of rewiring, STDP, and noise in synaptic weights. The Fokker-Planck equation provides a transparent link between local stochastic rules for synaptic plasticity and spine dynamics, and the resulting stationary distribution *p^*^*(**θ**) of network configurations **θ**. Learning is viewed from this perspective as convergence to a lower dimensional manifold of network configurations that provides good compromises between computational function and structural constraints. Structural constraints take the form of a prior in this model (see Fig. 4). One interesting benefit of this conceptual alternative to maximum likelihood learning is that the network immediately and automatically compensates for internal or external changes that modify the posterior distribution of network configurations (see Fig. 5 of [80]). But the underlying stochastic theory suggests that network configurations are likely to change continuously in functionally irrelevant dimensions – even in the absence of major perturbations.

A rethinking of the way in which network organization and plasticity is genetically encoded and implemented in the brain has been suggested by [82]. This challenge was motivated by the observation that the same neural circuit attains at different times and in different individuals the same performance with quite different parameter settings. The synaptic sampling perspective suggests an explanation for this observation: Each measurement of network parameters and synaptic connectivity provides a snapshot from an ongoing stochastic process.

**Figure 4:**
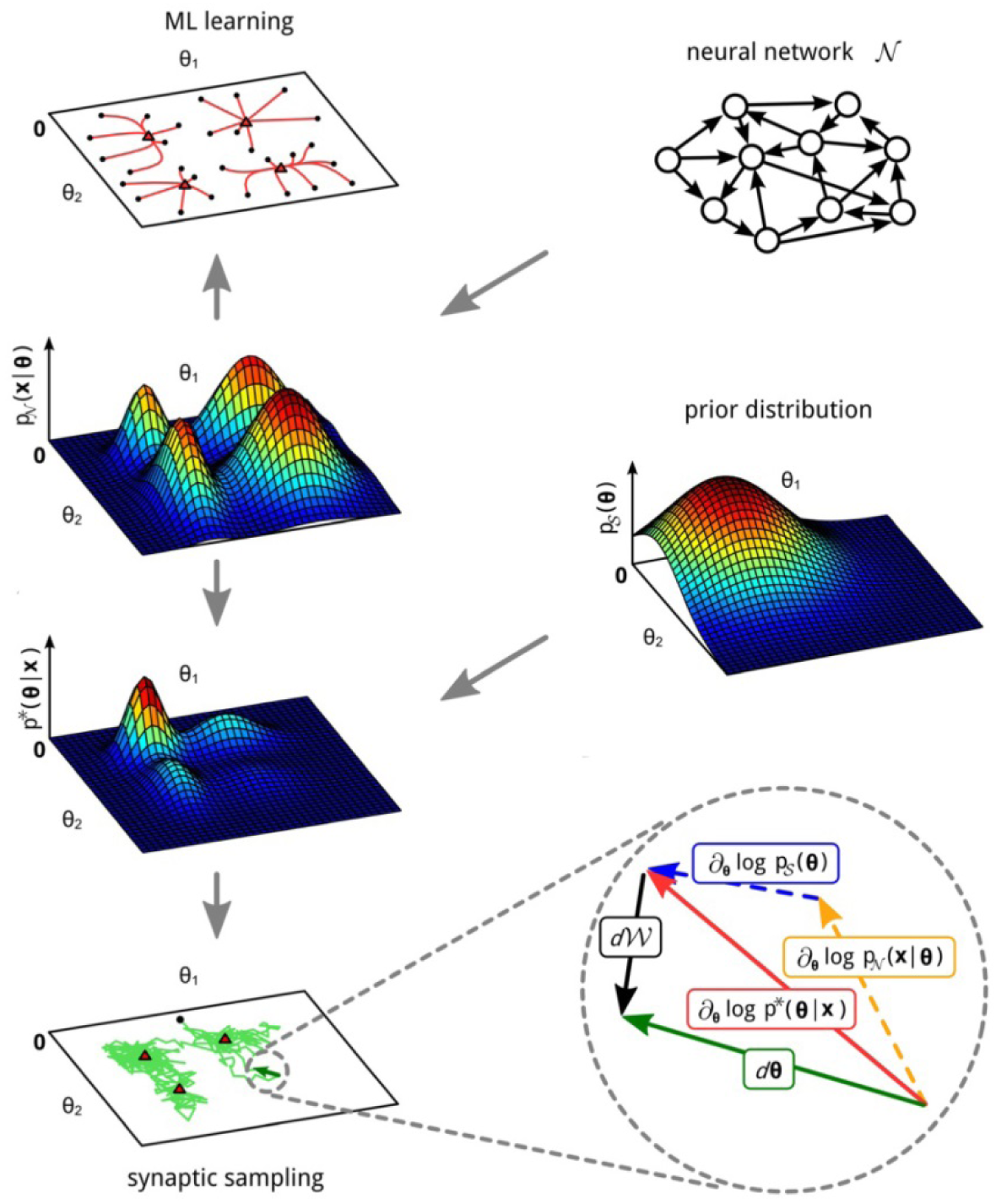
Two different options for the organization of network learning. Assume that some recurrent neural network *N* is given (top right), together with a generative model *p*N**(*x*|***θ***) (second plot from the top in the left column) for a given ensemble ***x*** of network inputs. Maximum likelihood learning moves the parameter vector ***θ*** of the network from a given initial state (black dot) to a local maximum of *p*N**(*x*|***θ***) (red triangles in top panel of left column). In contrast, the Bayesian synaptic sampling approach takes in addition a prior *p_s_* (***θ***) (middle panel in right column) into account – that could encode for example sparsity constraints – and aims at sampling network parameters *θ* from the posterior distribution *p^*^*(***θ***|***x***) α *p_s_*(***θ***) *p*N**(*x*|***θ***) (left column, 3^rd^ panel from the top). This can be achieved through a synaptic plasticity rule that takes the form of a stochastic differential equation with a drift term ∂***_θ_***log*p^*^*(***θ***| *x*) (red arrow in panel at the right bottom) that results from derivations of the log of the prior (blue arrow) and likelihood (yellow arrow), together with a stochastic diffusion term *dW* (black arrow). The Fokker-Planck equation implies that *p^*^*(***θ***| *x*) is the unique stationary distribution of this stochastic parameter dynamics (“synaptic sampling”). A sample trajectory of the parameter vector ***θ*** is plotted in green in the bottom left panel. Because of its stochastic component *dW* this learning approach can easily integrate stochastic spine dynamics with STDP, see (Kappel et al., 2015) [80] for details. The high-dimensional space of network parameters ***θ*** is replaced in this figure for illustration purposes by a 2D space. Fig. 4 is reprinted from [80] “Network plasticity as Bayesian inference” published by “PLOS Computational Biology, 11(11):e1004485, 2015”, with kind permission from The PLOS Journals.

## Conclusions

The four constraints/principles for models of brain computation and learning that I have discussed are compatible with each other. They have to be compatible, since experimental data tell us that they are all present in brain networks. But obviously there are tradeoffs between these principles. For example, more stereotypical network responses (principle 2) reduce the fading memory and kernel function (principle 1), see Fig. 12 in [50]. Hence I propose that the expression of each principle is regulated by the brain for each area and developmental stage in a task dependent manner.

Altogether I have argued that the currently avaible experimental data provide useful guidance for understanding how cognition and behaviour is implemented and regulated by networks of neurons in the brain. [83] had proposed to distinguish three levels of models for brain computations:

- the computational (behavioural) level
- the algorithmic level
- the biological implementation level.

Whereas substantial work had focused on the interconnection of these three levels from the top down, more detailed data on the biological implementation level provide now also a basis for creating bottom-up connections. We have seen that each of the four constraints from the biological implementation level has significant implications for models on the algorithmic and computational level.

## Acknowledgements

I would like to thank David Kappel, Robert Legenstein, and Zhaofei Yu for helpful comments, and Jonathan Wallis for sharing unpublished experimental data. This review was written under partial support the by the European Union project #604102.

